# Role of collagen-like protein BclA3 in the assembly of the exosporium layer of *Clostridioides difficile* spores

**DOI:** 10.1101/2021.06.21.449304

**Authors:** Marjorie Pizarro-Guajardo, Francisca Cid-Rojas, César Ortega-Lizárraga, Ana Inostroza, Daniel Paredes-Sabja

## Abstract

Newly formed spores are essential for persistence of *C. difficile* in the host, transmission to a new susceptible host (Deakin *et al.*, 2012b) and recurrence of CDI. BclA3 and BclA2 Spore surface proteins are expressed during sporulation under the control of mother-cell specific sigma factors of the RNA polymerase, SigE and SigK. Deletion of *bclA3* leads to spores with an electron-dense exosporium layer that lacks bump-like structures in the electron-dense layer and hair-like projections, both structures typically found in the wild type spore. Therefore, in this work, we have addressed the role of the exosporium collagen-like BclA3 glycoprotein in the assembly of the exosporium layer. Immunogold labelling of BclA2_CTD_ and BclA3_CTD_ indicates that both proteins are located in the hairs, with BclA2 located outermost of BclA3. Absence of BclA3 leads to spores with no hair-like projections, and absence of bumps in thick exosporium spores, a phenotype also expressed in by the deletion of the collagen-like region of BclA3. Overall, these results provide insights into the role of BclA3 in the assembly of the exosporium layer of *C. difficile* spores.

## Introduction

*Clostridioides difficile* is the leading cause of nosocomial antibiotic-associated diarrhea (Martin *et al.*, 2016, Rupnik *et al.*, 2009). Newly formed spores are essential for persistence of *C. difficile* in the host, transmission to a new susceptible host (Deakin *et al.*, 2012b) and recurrence of CDI, which is a major problem with an incidence of 25 to 65% of patients, associated to increase severity of symptoms and time in hospital settings (Lessa *et al.*, 2015). The surface of *C. difficile* spores play an important role in host-spore interaction (Mora-Uribe *et al.*, 2016, Calderón-Romero *et al.*, 2018, Paredes-Sabja & Sarker, 2012). In the spore of epidemic *C. difficile* strains, such as RT27 and RT78, the most external layer, the exosporium, is surrounded by an ultrastructure defined as hair-like projection, which appears as a layer of ∼80 nm of disheveled hairs, differently to the hair-like nap in *B. anthracis* spore, where all hairs are extended to the exterior (Thompson *et al.*, 2011). However, hair-like extensions are absent in 630 strain spores (Paredes-Sabja *et al.*, 2014). The role of this structure in pathogenesis is unknown.

Spore surface proteins are expressed during sporulation under the control of mother-cell specific sigma factors of the RNA polymerase, SigE and SigK (Fimlaid *et al.*, 2013). By sonication of *C. difficile* spores, trypsin digestion of extracted proteins and MS/MS, spore coat and exosporium proteins have been identified (Díaz-González *et al.*, 2015). Some of the surface spore proteins are collagen-like proteins, characterized by a large inner domain with several GXY repeated pattern, flanked by globular NTD and CTD (Díaz-González *et al.*, 2015, Pizarro-Guajardo *et al.*, 2014). Collagen-like proteins are present in several spore structures, such as appendages in *Clostridium teaniosporium* (Steichen *et al.*, 2003) and external nap in *Bacillus anthracis* (Steichen *et al.*, 2003), where collagen-like proteins tend to form trimers by the globular carboxi-terminal domain (CTD) that allow the formation of triple helical collagen-like structures (Steichen *et al.*, 2003, Yu *et al.*, 2014).

In *C. diffi*cile genome, three homologues of the *bclA* loci are present in most *C. difficile* strains (Pizarro-Guajardo *et al.*, 2014). However, most epidemically-relevant strains belonging to ribotype 027 have only full-length *bclA2* and *bclA3* open reading frames, while a nonsense mutation allows the expression of a pseudogenized *bclA1* (Pizarro-Guajardo *et al.*, 2014, Phetcharaburanin *et al.*, 2014, Hosseini, 2018). A previous study demonstrated that insertional inactivation of *bclA1* and *bclA2* genes lead to spores with a defectively assembled electron-dense exosporium layer in strain 630, but normal morphology for the mutant in *bclA3* (Phetcharaburanin *et al.*, 2014). For epidemic strain R20291, our results indicate that a deletion of *bclA3* leads to spores with an electron-dense exosporium layer that lacks bump-like structures in the electron-dense layer and hair-like projections, both structures typically found in the wild type spore (Castro-Córdova *et al.*, 2020). The gene *bclA3* is the second element in a bicistronic operon, where the first cistron, *sgtA*, has been characterized as a glycosyltransferase enzyme that is required for the glycosylation of collagen-like fragments of BclA3 (Aubry *et al.*, 2020, Strong *et al.*, 2014).

Therefore, in this work, we have addressed the role of the exosporium collagen-like BclA3 glycoprotein in the assembly of the exosporium layer. Immunogold labelling of BclA2_CTD_ and BclA3_CTD_ indicates that both proteins are located in the hairs, with BclA2 located outermost of BclA3. Absence of BclA3 leads to spores with no hair-like projections, and absence of bumps in thick exosporium spores, a phenotype also expressed in by the deletion of the collagen-like region of BclA3. Overall, these results provide insights into the role of BclA3 in the assembly of the exosporium layer of *C. difficile* spores.

## Materials and Methods

### Bacterial strains and growth conditions

*C. difficile* strains (see Supplementary Table 1) were routinely grown at 37 °C under anaerobic conditions in a Bactron III-2 anaerobic chamber (Shellab, USA) in BHIS medium: 3.7% weight vol−1 brain heart infusion broth (BD, USA) supplemented with 0.5% weight vol−1 yeast extract (BD, USA) and 0.1% weight vol−1 L-cysteine (Merck, USA) or on BHIS agar plates. Escherichia coli strains were routinely grown aerobically at 37 °C under aerobic conditions with shaking at 1 × g in Luria-Bertani medium (BD, USA), supplemented with 25 μg mL−1 chloramphenicol (Merck, USA), where appropriate.

### *C. difficile* mutant construction of mutant strain in the CLR domain in *bclA3* gene (*bclA3*_ΔCLR_)

In order to construct a R20291 *C. difficile* strains which lacks the CLR domain in *bclA3* gene (*bclA3*_ΔCLR_), *C. difficile* genome FN545816 from the EMBL/GenBank databases was employed for primer design. In-frame deletions were made by allelic exchange using *pyrE* alleles (Ng *et al.*, 2013). To remove the CLR segment from *bclA3* locus, left homology arm was constructed by amplification of 1676 bp fragment including 1529 nt upstream *bclA3* start codon, first 138 nt which codify 46 aa corresponding to amino-terminal domain (NTD), plus 9 nt corresponding to the first GXX of collagen-like region, flanked by 5’ SbfI and 3’ BamHI restriction sites, which was cloned in those restriction sites into plasmid pMTL-YN4. For right homology arm, a 1042 nt fragment, starting at 1516 nt downstream start codon, was amplified with 5’ BamHI and 3’ AscI restriction sites, and cloned in those restriction sites into pMTL-YN4 containing left homology arm. To verify the correct construction of the plasmids, all constructs were Sanger sequenced.

The plasmid obtained were transformed into *E. coli* CA434 and mated with *C. difficile* R20291 Δ*pyrE* (Ng *et al.*, 2013). *C. difficile* transconjugants were selected by sub-culturing on BHIS agar containing 15 µg/mL thiamphenicol (Sigma-Aldrich USA), 8 µg/mL cefoxitin (Sigma-Aldrich USA) and 250 µg/mL cefoxitin, and re-streaked five times. In order to select for plasmid excision, the single-crossover mutants identified were streaked onto CDMM (Karasawa *et al.*, 1995) with 1.5% agar supplemented with 2 mg/mL 5-Fluoroorotic acid (US Biological, USA) and 5 µg/mL uracil (Sigma-Aldrich USA). Confirmation of plasmid excision was made by negative selection in BHIS-15% thiamphenicol plates. The isolated FOA-resistant and thiamphenicol sensitive colonies were screened using the primer pair P664 (FP-bclA3-detect) / P665 (RP-bclA3-detect) for the *bclA3* mutant. The mutant was whole genome sequenced to confirm the genetic background and that no additional SNPs were introduced during the genetic manipulation. For correction of the *pyrE* mutation, transconjugants with pMTL-YN2C were streaked onto minimal media without uracil or FOA supplementation, and developed colonies were analysed further. To verify correct insertion, genomic DNA was extracted and sequenced.

### Spore purification of *C. difficile* R20291

*C. difficile* strains used in this study are listed in Table 1. To produce pure spores, *C. difficile* was routinely grown under anaerobic conditions on BHIS broth (3.7% Brain Heart Infusion supplemented with 0.5% yeast extract, 1% cysteine). 16 h cultures were diluted to 1:500 dilution and 100 ul were plated onto 70:30 (63 g Bacto peptone, 3.5 g protease peptone, 11.1 g BHI medium, 1.5 g yeast extract, 1.06 g Tris, 0.7 g NH_4_SO_4_, 15 g agar per litter) or TY (3% Trypticase Soy-0.5% yeast extract) agar plate and incubated at 37 °C for 7 days. After incubation, plates were scraped up with 1 ml of ice-cold sterile water. Spores were gently washed five times (resuspension in ice-cold sterile water and centrifugation at 16,000 × g for 5 min). Next, spores were gently loaded onto a 1.5 ml tube containing 300 μl of 45% Nycodenz solution and centrifuged for 40 min at 16,000 × g. After centrifugation, the supernatant was removed, and the spore pellet was washed five times (Calderón-Romero *et al.*, 2018). The spores were counted in a Neubauer chamber and concentration adjusted at 5 × 10^9^ spores/ml to store at -80°C until use.

### Transmission electronic microscopy (TEM)

Spores (2×10^8^) were fixed with 3% glutaraldehyde on 0.1 M cacodylate buffer (pH 7.2), incubated overnight at 4°C, and stained for 30 min with 1% tannic acid. Samples were further embedded in a Spurr resin (Paredes-Sabja *et al.*, 2012). Thin sections of 90 nm were obtained with a microtome, placed on glow discharge carbon-coated grids and double lead stained with 2% uranyl acetate and lead citrate. Spores were analyzed with a Philips Tecnai 12 Bio Twin microscope at Unidad de Microscopía Avanzada at the Pontificia Universidad Católica de Chile. ImageJ software was employed to calculate thickness of spore layers.

### Immunogold

To locate BclA2 and BclA3 in the spore, 4×10^7^ R20291 spores were incubated in PBS–0.2% BSA containing 1:200 mouse serum from immunized animal with BclA2_CTD_ and BclA3CTD, respectively, for 1 hr at 37 ºC. The excess of antibody was eliminated by three cycles of centrifugation at 13,000□J×□Jg for 5□Jmin and resuspension in PBS–0.1% BSA. Spore suspensions were then incubated for 1□Jh with 1:20 donkey anti-mouse IgG antibody coupled to 12-nm gold particles (Sigma G7777) in PBS–1% BSA for 1□Jhr at RT, and were washed by centrifugation at 13,000□J×□Jg for 5□Jmin. Subsequently, samples were fixed and processed, as described. ImageJ software was employed to calculate distance from spore surface to gold label.

## Results

### Revisiting *C. difficile* spore exosporium thickness

As previously described, *C. difficile* sporulation process generates two distinctive exosporium morphotypes, thick and thin exosporium (Pizarro-Guajardo *et al.*, 2016a, Pizarro-Guajardo *et al.*, 2016b, Paredes-Sabja *et al.*, 2014), which differs in the electron-dense layer; while a ∼20 nm electron-dense layer surrounds the coat spore in thin exosporium spore, the electron-dense layer is made of bumps that can reach ∼100 nm. Despite this difference, both morphotypes share the hair-like projection ultrastructure (Pizarro-Guajardo *et al.*, 2016b, Pizarro-Guajardo *et al.*, 2016a). It remains unclear if the site of anchorage of these filaments is the coat with the electron-dense bumps being formed surrounding these filaments, or otherwise, the filaments are located on the surface of the bumps, in the thick exosporium spore. Here, we analyzed a total of 39 thin exosporium and 35 thick exosporium spores from at least three independent spore preps. Measure included a tangential line over the inner membrane layer, where a line projected in 90º was used to guide the angle of the measures (Figure 1A). Thick exosporium spores were measured in the maximum thickness of the exosporium (Figure 1A, dotted line). The electron dense layer is thicker in thick exosporium spores, with significant difference (p<0.0001; unpaired t-test). Distribution of thickness follows a bimodal distribution of thick and thin-exosporium spores (Figure 1D).

**Figure 1.**
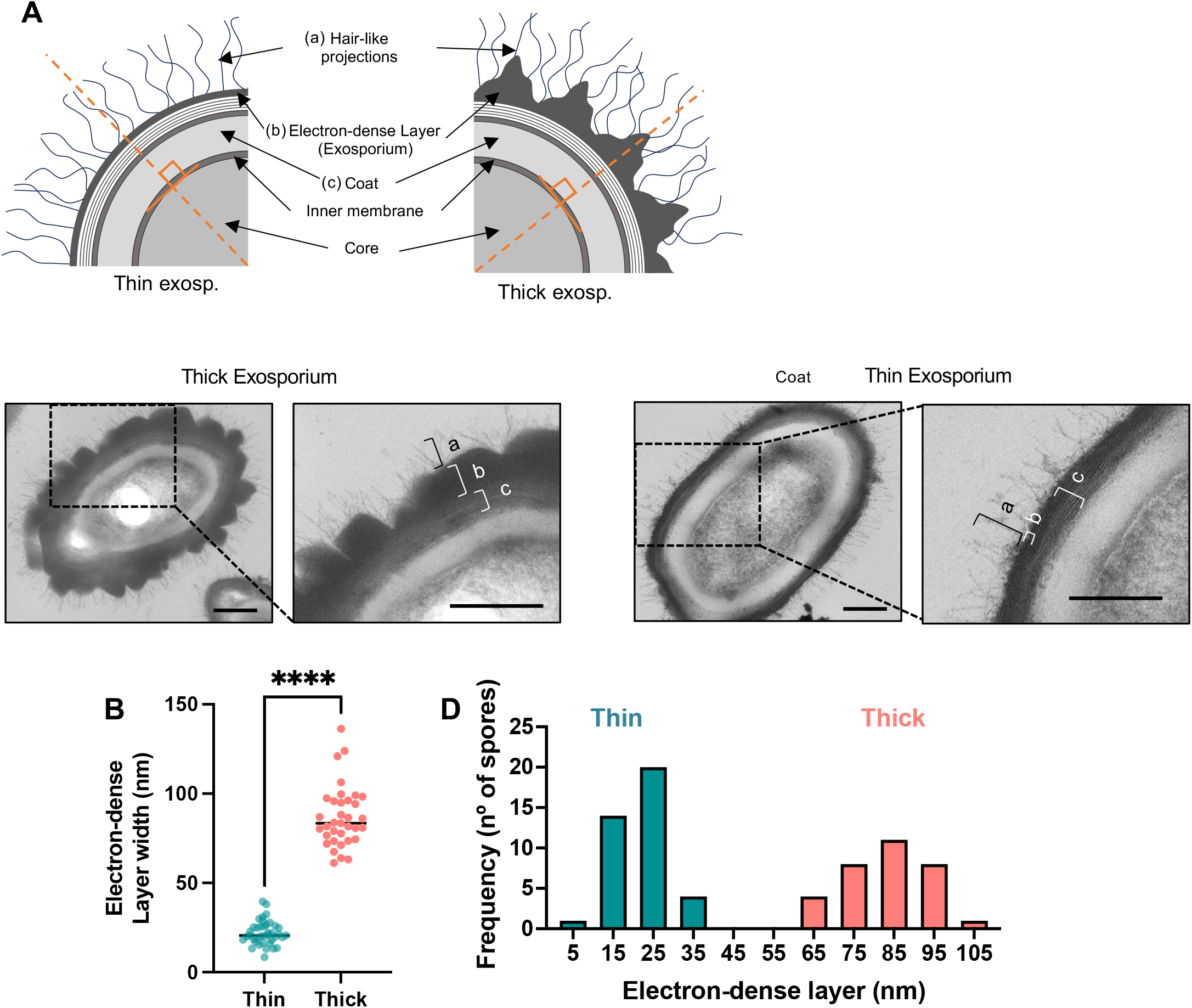
Bimodal-exosporium morphotypes. A) Schematic representation of thick and thick exosporium spores and representative transmission electro micrographs of *C. difficile* spores of strain R20291 with a thin-or a thick-exosporium morphotype. B) Measure of the width of electron-dense layer in thin and Thick exosporium spores (Mann Whitney test); ****, *p* < 0.0001. C) Frequency distribution of thickness follows a bimodal distribution of thick and thin-exosporium spores.

### Collagen-like proteins BclA2 and BclA3 are part of the hair-like projections of *C. difficile* spores

R20291 strain encodes two collagen-like proteins, BclA2 and BclA3 (Pizarro-Guajardo *et al.*, 2014). These proteins have a central collagen-like region, a similar size CTD, and a different size NTD (i.e. BclA2 has a shorter NTD compared to BclA3, 5 versus 46 amino acid-length) (Figure 2a). We have previously shown that in strain 630, which has a functional BclA1 protein, the NTD seems to be oriented in an inward orientation, with the CTD oriented outwards (Pizarro-Guajardo *et al.*, 2014). In addition, transmission electron micrographs of *bclA3* mutant spores demonstrates that BclA3 is required for the formation of the hair-like projections (Castro-Córdova *et al.*, 2021); however, it is unclear whether these hair-like projections are formed by BclA2 and/or BclA3.

**Figure 2.**
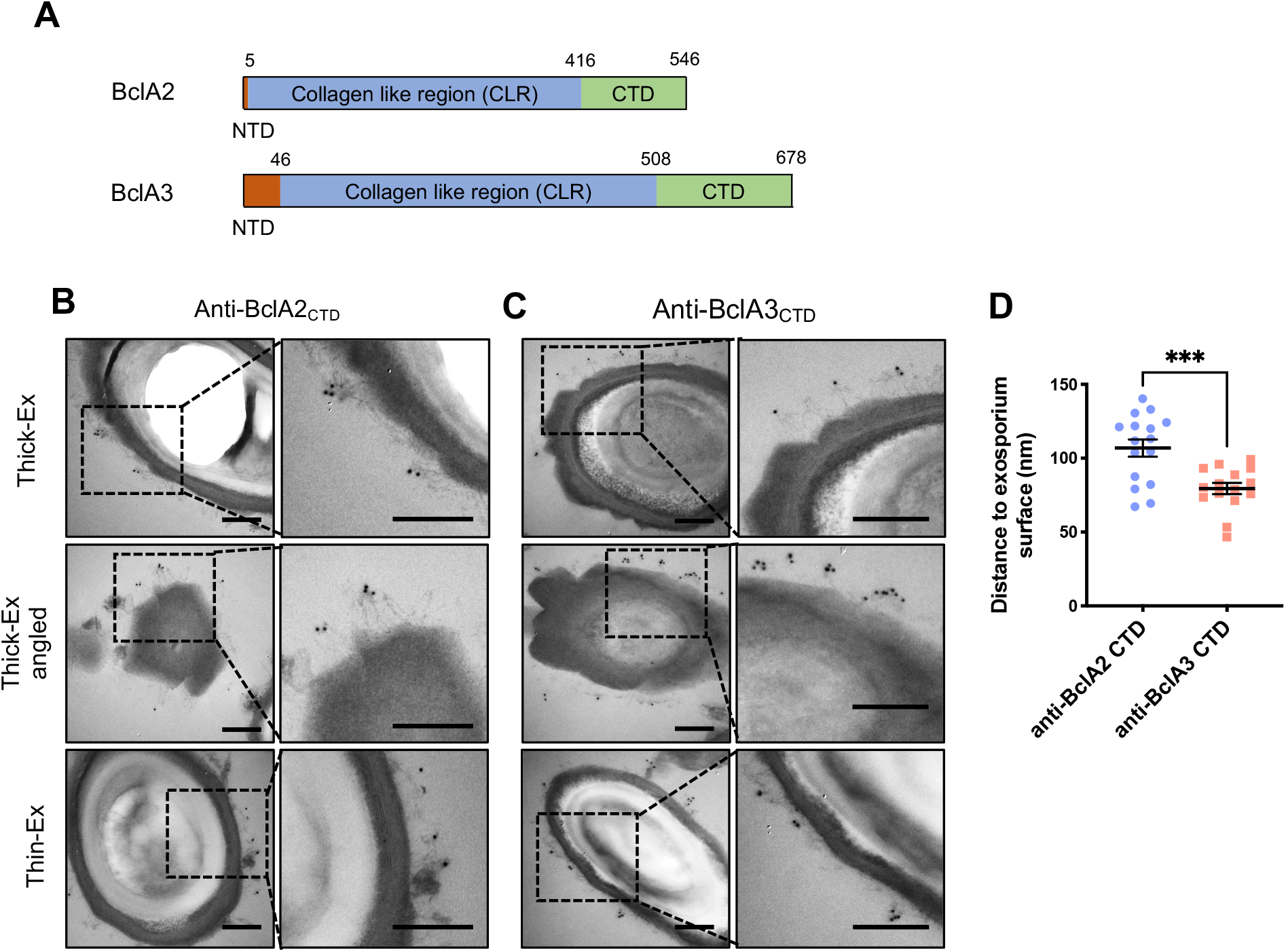
Immunogold labeling of hair-like extensions trough BclA2 and BclA3 targeting in R20291 spores. A) Schematic representation of domains in BclA2 and BclA3. Red: NTD, green: CTD, yellow: collagen like repeats domain. Numbers indicate the aminoacidic position where domains end. B) Targeting of hair-like extensions with anti-BclA2 antibody and C) anti-BclA3 antibody, in wild type R20291. Right panel is a zoom of left panel. Bar scale: 200 nm. Black squared dot shows the zoomed area. D) Distance from the surface of the spore to the gold label. Every dot represent one spore, as the average distance of 1 to 9 gold particles to the electron-dense surface. Statistical test: Unpaired t-test.

To solve this question, we employed antibodies against BclA2_CTD_ and BclA3_CTD_ to tag these proteins in *C. difficile* spore in an immunogold assay. Immunoelectron micrographs demonstrate that both antibodies labelled the tip of the hair-like projections in both exosporium morphotypes (thick and thin exosporium) (Figure 2B and Figure 2C), indicating that BclA2 and BclA3 are part of the hair-like projections, and that both CTD are oriented outwards. Next, we quantified the distance of the labelled CTD of BclA2 and BclA3 to the electron dense surface of spores, which indicates that BclA2 could have a location 25 nm outer that BclA3 (Figure 2D). This outer position of BclA2_CTD_ respect to BclA3_CTD_ is independent of the exosporium type.

### Thick and thin-exosporium layer *C. difficile* spores lack the hair-like projections in the absence of BclA3

Our previous work demonstrated that inactivation of BclA3 lead to the absence of hair-like projections. While in the wild type strain, hair-like projections and exosporium electron-dense bumps were present in thick and thin exosporium spores; as previously described (Castro-Córdova *et al.*, 2020), Δ*bclA3* spores exhibited no hair-like projections in thick and thin exosporium spores (Figure 3A). Notably, we identified in thick exosporium spores a smooth electron-dense layer without bumps (Figure 3A). Upon complementation of the Δ*bclA3* mutant strain with *bclA3* into the *pyrE* loci, we observed that spores of the Δ*bclA3/bclA3+* strain restored the hair-like projections and electron dense bump formation (Figure 3A, B), demonstrating that BclA3 is required for the formation of the hair-like projections and electron-dense bumps of the exosporium layer. Upon analysis of the spore-coat thickness, we observed that BclA3 had no impact in the thickness of the spore-coat (Figure 3A and 3B). Collectively, these results demonstrate that the role of BclA3 in external layers morphology is restricted to the morphology of the exosporium, due to the absence of alterations in spore coat layer. Specifically, BclA3 plays a key morphological role in the formation of the electron-dense bumps and hair-like projections.

**Figure 3.**
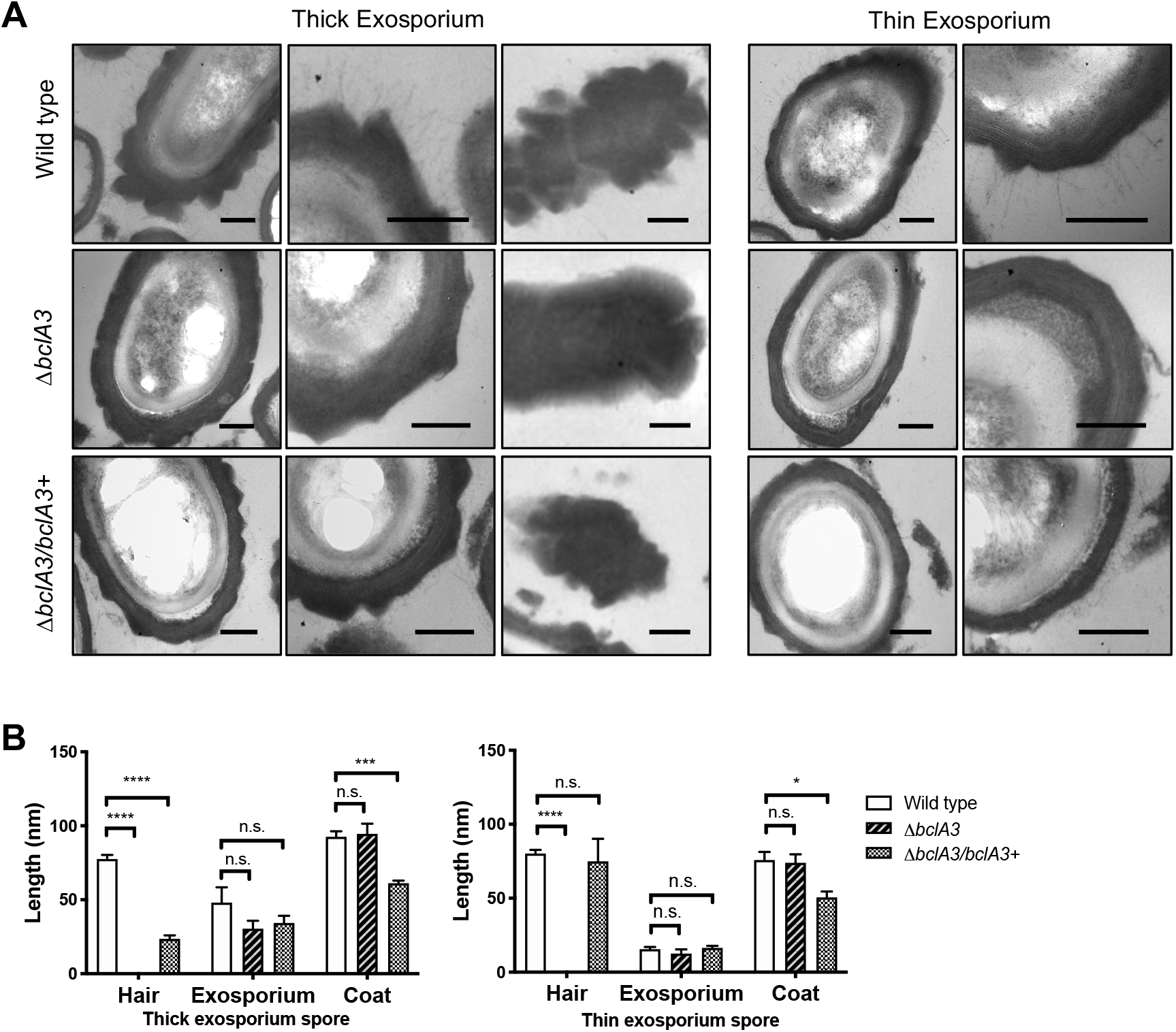
BclA3 have a role in hair-like ultrastructure in R20291 spores. A) Purified spores from R20291, *ΔbclA3* and complemented. *C. difficile* strains were fixed and analyzed by transmission electron microscopy. Spores from wild type, Δ*bclA3* and Δ*bclA3/bclA3*+ were classified as thick or thin exosporium. Scale bar: 200nm. B) Thickness of each layer (hair, exosporium and coat) was measured at 6 different points (n = 7; 6 measures per spore). 2-way ANOVA, Turkey’s multiple comparison test.

### Lack of BclA3 leads to enrichment of *C. difficile* spores with appendages

Another feature of *C. difficile* spores that has been recently described is the presence of appendages in the spore poles (Pizarro-Guajardo *et al.*, 2020, Antunes *et al.*, 2018). Appendages are defined as a prolongation of the exosporium in one polar region of the spore (Pizarro-Guajardo *et al.*, 2020, Antunes *et al.*, 2018), which is dependent on CdeM and is related to faster germination (Antunes *et al.*, 2018). In order to evaluate impact of BclA3 in the appendage frequency, we used phase contrast microscopy to characterize populations of wild type and *ΔbclA3* mutant spores. Phase contrast microscopy shows polar appendage in two different morphologies: appendage type 1, a polar prolongation, and appendage type 2, diffuse and round prolongation (Figure 4A). In the wild type strain, most of the spores (84%) had no appendage, while the rest of the spores were distributed between App-1 (8.9%) and App-2 (6.4%). In contrast, for the *ΔbclA3* mutant, the spores lacking appendage decreased to 59%, while spores with App-1 and App-2 increased to 18% and 23%, respectively (Figure 4B). Notably, restoration of *bclA3* restores the spore proportions of appendages to similarly levels as wild type strain (Figure 4B), indicating that presence of BclA3 in the exosporium spore favors the spore type without appendage.

**Figure 4.**
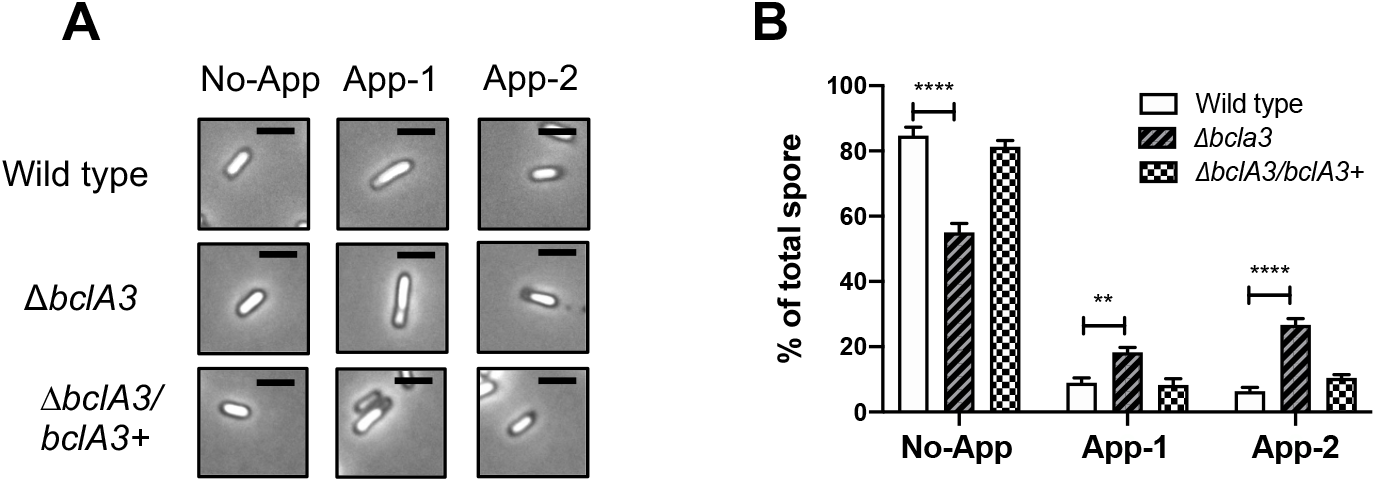
Appendage occurrence in R20291 and bclA3 mutant spores. A) Example of phase-contrast microscopy spores showing no-appendage (No-app), appendage type 1 (App-1) and appendage type 2 (App-2). Scale bar: 1 μm. B) Percentage of appendage type. N = 727 wild type, 985 *bclA3-*, 832 *bclA3-/bclA3*+. 2-way ANOVA, Multiple comparison test.

### The collagen-like region of BclA3 is required for the formation of the hair-like projections and electron-dense bumps of the exosporium layer

Work in *Bacillus anthracis* has demonstrated that the length of the hairs of the exosporium layers depended on the collagen-like region (CLR) of the BclA glycoprotein (Sylvestre *et al.*, 2003). Since our results demonstrate that the hair-like projections are formed by BclA2 and BclA3, but *bclA3* is essential for the formation of the hair-like projections, we asked if the CLR of *C. difficile* BclA3 is required for the formation of these projections. To assess this, we constructed a *C. difficile* strain with a deletion in the CLR region of the *blcA3* gene (*bclA3*_ΔCLR141-1509_) (Figure 5A). Transmission electron micrographs demonstrate that while long hair-like projections are exhibited in wild type spores, mutant strain lacking BclA3 CLR domain did not exhibit hair-like projections or electron-dense bumps in the exosporium layer; *bclA3*_ΔCLR141-1509_ spores resembled Δ*bclA3* spore ultrastructural phenotype (Figure 5B). These results demonstrate that the CLR of BclA3 is essential for the proper formation of the hair-like projection structures and bump formation in the exosporium of *C. difficile* spores.

**Figure 5.**
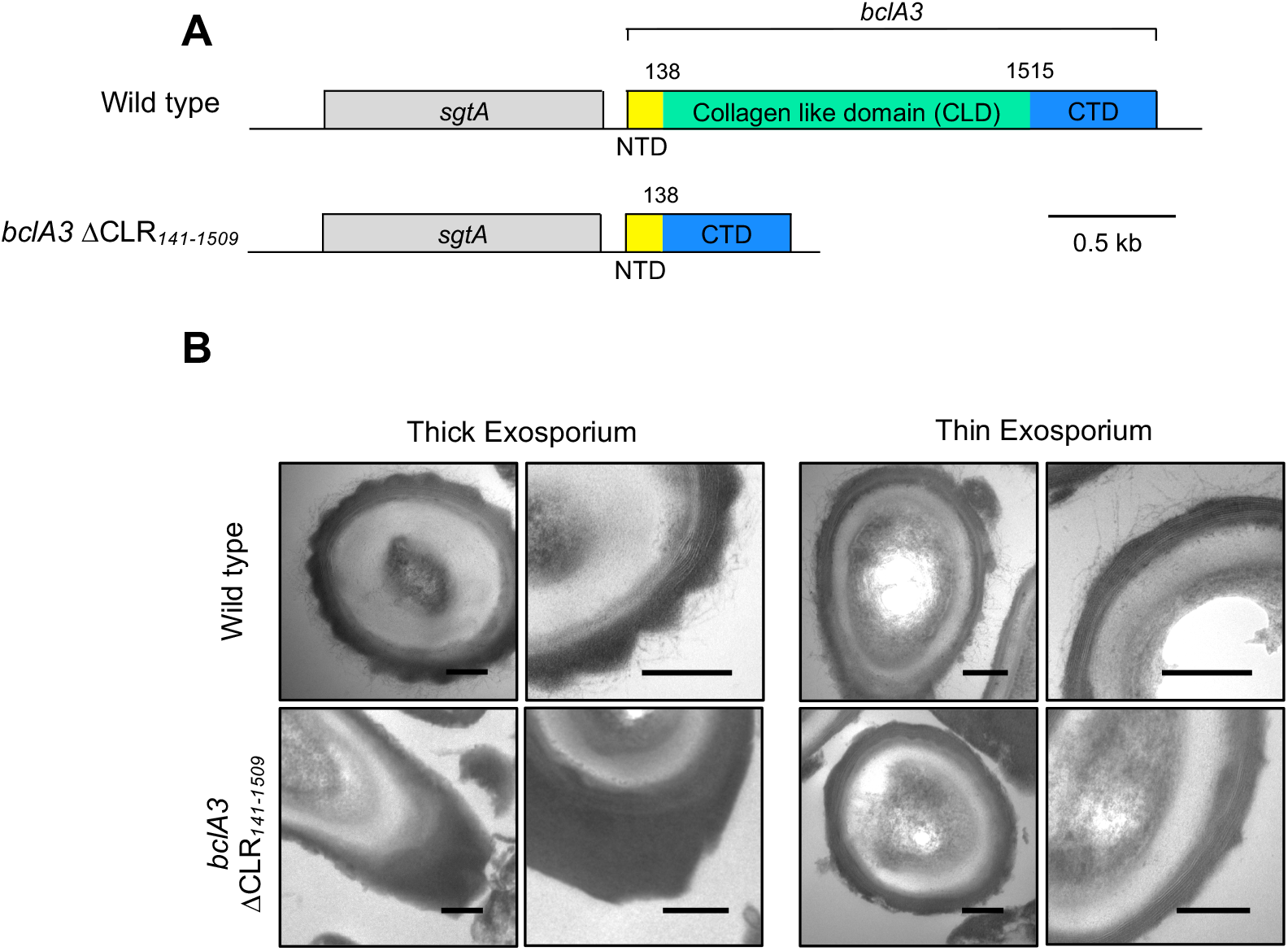
Lack of collagen-like domain in bclA3 gene impairs the hair formation and impacts exosporium morphology. A) Schematic representation of reverse R20291 DNA strand in wild type strain and mutant in CLR portion of *bclA3* gene (*bclA3* ΔCLR_*142-1509*_) in reverse strand. B) TEM imagen of thick and thin exosporium spores of *bclA3* ΔCLR_*142-1509*_, showing hair-exosporium lack and lack of bumps. Scale bar: 200 nm.

## Discussion

During CDI, *C. difficile* spore formation is essential in the recurrence of the disease (Deakin *et al.*, 2012a), and the underlying mechanisms that correlate *C. difficile* spore persistence and recurrence of the disease stared to be unveiled. In this sense, the outermost layer of the spore, the exosporium, is critical to understand CDI pathogenesis and developing of novel therapies. Recently, a novel mechanism for interaction between the spore and the intestinal mucosa has been described, in which BclA3 is essential for the formation of the hair-like projections, but we know little of the role in exosporium assembly and spore properties (Castro-Córdova *et al.*, 2020). Therefore, BclA3 has major contribution in *C. difficile* persistence and initiation of recurrence, that suggest that blocking its interactions with the host environment could improve the clinical outcome of recurrence. In this work, we further dissected the morphogenic role of collagen-like proteins BclA3 in the exosporium of *C. difficile* spore. Previous work on the role of BclA3 in *C. difficile* spore assembly made on the strain 630 cannot be complementary with these results, due to the lack of hairs in 630 strain.

The first major finding of this work is the labeling of BclA2_CTD_ and BclA3_CTD_ at the tip of the hair-like extension, that indicates that hair-like extensions are made by both, BclA2 and BclA3. A mutant Δ*bclA3* strain spore has no capability to form hair-like extensions (Castro-Córdova *et al.*, 2020), indicating that the presence of only BclA2 is not sufficient to build the hair-like projection structure. However, it is described that BclA2 can form collagen-like structures susceptible to digestion by collagenase (Pizarro-Guajardo *et al.*, 2014). As only BclA3 is essential for hair-like projection structure, and BclA2_CTD_ was tagged in the structure, it suggests that BclA2 is capable to adhere to hair-like structures once they are formed by BclA3.

Whether BclA2 is forming collagen-like structure with BclA3 after BclA3 spore assembly remains unknown. The location of BclA2_CTD_ more external than BclA3_CTD_ suggest that BclA3 is necessary to assemble of BclA2 in the collagen structure, which is supported by the lack of hair when BclA2 is transcribed but not *bclA3* in the mutant Δ*bclA3*. Detection of BclA2_CTD_ in the spore Δ*bclA3* suggest that BclA2 can assemble in the spore independently of BclA3, but in this assemble is not capable of structure hairs.

Bump formation in exosporium seems to be a consequence of hair structure, since all mutations affecting BclA3 (Δ*bclA3*, lack of collagen-like repeat domain) impair the hair formation, but also prevent the bumps structure to form in the exosporium. It is interesting to observe that mutant strain lacking the collagen rich region of BclA3, the CLR, lacks the hair-like extension but the exosporium shows a reduced bump exosporium, which suggest that a slightly development of exosporium bump, but not all, is developed in the presence of BclA3_NTD-CTD_ only. It is described that only NTD of BclA3 and BclA2 are sufficient to anchor the protein to the spore (Pizarro-Guajardo *et al.*, 2014), which suggest that CTD domain of both proteins is exposed to the surface, as confirmed by immunogold experiments in this work.

## 5. Acknowledgements

This work was supported by FONDECYT Regular 1191601, FONDECYT Regular 1151025, and Millennium Science Initiative Program–NCN17_093, and Fondo de Fomento al Desarrollo Científico y Tecnológico (FONDEF) ID18|10230 to M.P-G and D.P-S. Support was also provided by start-up funds from Texas A&M University to D.P-S.

## Tables

**Table S1. Primers used in this work to create SNAP fusions.**

**Table S2. Plasmids and strains used in this work**

